# The Deubiquitinating Enzyme USP37 Stabilizes CHK1 to Promote the Cellular Response to Replication Stress

**DOI:** 10.1101/652891

**Authors:** Mayank Singh, Amy C. Burrows, Adrian E. Torres, Debjani Pal, Benjamin Stromberg, Christine Insinna, Andrew Dickson, Christopher J. Westlake, Matthew K. Summers

**Affiliations:** Department of Radiation Oncology, The Ohio State University Wexner Medical Center and Comprehensive Cancer Center, Columbus OH, 44120 USA; Department of Cancer Biology, Lerner Research Institute, Cleveland Clinic, Cleveland OH, 44195 USA; NCI-Frederick National Laboratory, Laboratory of Cellular and Developmental Signaling, Frederick, MD 21702, USA; Department of Medical Oncology (lab) BRAIRCH AIIMS Delhi, Delhi India 110029

**Keywords:** cell cycle, CHK1, replication checkpoint, USP37

## Abstract

The deubiquitinating enzyme USP37 is known to contribute to timely S-phase onset. However, it is not clear if USP37 is essential beyond S-phase entry despite expression and activity of USP37 peaking within S-phase. Here, we determine that USP37-depleted cells exhibited altered S-phase kinetics and harbored increased levels of the replication stress and DNA damage markers γH2AX and 53BP1 in response to perturbed replication. Depletion of USP37 also reduced cellular proliferation and led to increased sensitivity to agents that induce replication stress. Underlying the increased sensitivity, we found that the checkpoint kinase CHK1 is destabilized in the absence of USP37, attenuating its function. We further demonstrated that USP37 deubquitinates and stabilizes CHK1. Our data contribute to an evolving understanding of UPS37 as a multifaceted regulator of genome regulation.

## INTRODUCTION

Cells encounter a multitude of intrinsic and extrinsic barriers to accurate DNA replication. To ensure that replication is error free, eukaryotes possess a conserved checkpoint that monitors the replication process. Upon replication stress (e.g., stalled replication fork, nucleotide deficiency, DNA damage), the ATR kinase is recruited to DNA (Cimprich and Cortez, 2008; Wang et al., 2006). The checkpoint mediators RAD17 and CLASPIN then recruit the effector kinase CHK1, which is phosphorylated and activated by ATR at S317 and S345. In turn, CHK1 phosphorylates a number of proteins to prevent origin firing and entry into mitosis as well as promoting DNA repair (Cimprich and Cortez, 2008; Merry et al., 2010). In the absence of CHK1 recruitment and activation upon replication stress, cells maintain high levels of CDK activity, continue to fire origins, and accumulate additional damage and chromosomal abnormalities. These cells are thus highly sensitive to additional replication stress. Importantly, high levels of replication stress are associated with high rates of proliferation during early development and expression of multiple oncogenes (e.g., Cyclin E, c-MYC) (Bartkova et al., 2006; Bester et al., 2011; Halazonetis et al., 2008; Hoglund et al., 2011; Murga et al., 2011; Schoppy et al., 2012). Transformation thus requires that cells survive with these abnormal levels of replication stress (Bartkova et al., 2006). As a result, transformed cells are highly dependent on the ATR-CLASPIN-CHK1 pathway for survival and are sensitive to agents that either induce additional stress or inhibit this critical checkpoint (Hoglund et al., 2011; Murga et al., 2011; Schoppy et al., 2012).

Intriguingly, premature CHK1 activation may drive S-phase entry and failure to down-regulate CHK1 activation is also detrimental to cellular survival. CHK1 activity is thus tightly regulated on multiple levels including ubiquitin-mediated degradation of key components of the replication stress response pathway. During G1, the anaphase promoting complex/cyclosome in complex with the activator CDH1 (APC/C^CDH1^) ubiquitin ligase prevents premature accumulation of both CLASPIN and RAD17 and hence premature S-phase entry (Bassermann et al., 2008; Faustrup et al., 2009; Gao et al., 2009). During G2 and mitosis, APC/C^CDH1^ and a second ubiquitin ligase, Skp1-Cul1-F-box in complex with the F-box and activator protein, βTRCP, (SCF^βTRCP^), prevent additional CHK1 activation by triggering the destruction of RAD17 and CLASPIN, respectively (Mailand et al., 2006; Mamely et al., 2006; Peschiaroli et al., 2006). Additionally, the E3 ligases, HERC2 and BRCA1, have also been demonstrated to ubiquitinate CLASPIN(Sato et al., 2012; Yuan et al., 2014; Zhu et al., 2014). Further underscoring the need for tight control of CHK1 activation, multiple deubiquitinating enzymes (DUBs), including USP7, USP9x, USP28, and USP29, enhance CHK1 activation by stabilizing CLASPIN, whereas several recent reports demonstrate that USP20 promotes CHK1 function by stabilizing both RAD17 and CLASPIN (Bassermann et al., 2008; Faustrup et al., 2009; Martin et al., 2015; McGarry et al., 2016; Shanmugam et al., 2014; Yuan et al., 2014; Zhang et al., 2006; Zhu et al., 2014). CHK1 itself is also targeted for destruction. Upon activation, CHK1 adopts an open conformation, which exposes degrons that are recognized by SCF^FBX6^ and the Cullin 4-based cullin ring ligase (CRL4^CDT2^) ubiquitin ligases (Huh and Piwnica-Worms, 2013; Leung-Pineda et al., 2009). Not surprisingly, DUB activity also plays a role in the maintenance of CHK1 levels. However, there are few examples of CHK1 stabilization by DUBs in comparison to their involvement stabilizing proteins that direct the activation of CHK1. To date, only USP7 and USP1 have been implicated in the maintenance of CHK1 levels and only USP7 has been demonstrated to act directly on CHK1 (Alonso-de Vega et al., 2014; Guervilly et al., 2011; Zhang et al., 2014). Together, these mechanisms confine checkpoint activity to S-phase, promote recovery from checkpoint activity and prevent hyperactivation of the checkpoint.

We recently determined that, similar to CLASPIN, the expression of the DUB USP37 is primarily confined to S-phase by the concerted efforts of APC/C^CDH1^ and SCF^βTRCP^, indicating roles of USP37 in the replication process (Burrows et al., 2012; Huang et al., 2011). Indeed, USP37 regulates S-phase entry via stabilization of Cyclin A and the licensing factor, CDT1 (Hernandez-Perez et al., 2016). In the absence of USP37 replication fork speed is also delayed (Hernandez-Perez et al., 2016). However, the function(s) of USP37 after S-phase entry and the biological consequences of deficiencies in USP37 function are not well characterized. Here, we describe a requirement for USP37 in the tolerance of replication stress, which is reflected in the ability of USP37 to deubiquitinate the active form of CHK1 leading to its stabilization and the maintenance of CHK1 activity.

## RESULTS

### USP37 Promotes Efficient Replication

We and others have previously demonstrated that USP37 regulates the timing of S-phase entry, at least in part by facilitating the stability and accumulation of Cyclin A and CDT1 during G1 (Hernandez-Perez et al., 2016; Huang et al., 2011). However, it remains unclear whether USP37 is required beyond S-phase entry. We next sought to determine whether loss of USP37 impacts S-phase progression.

Because nocodazole-synchronized USP37-depleted cells enter S-phase with delayed kinetics we sought to analyze the effects of USP37-depletion in otherwise unperturbed cells that had already entered S-phase. We thus labeled asynchronous populations of control or USP37-depleted HCT116 cells with EdU, to mark replicating cells, and analyzed these cells by flow cytometry (Figures 1A-B). Consistent with a recent report, USP37-depletion had no gross impact on the overall cell cycle profile of asynchronous populations (Figure 1A, top panels) (Hernandez-Perez et al., 2016). We then examined cell cycle progression by asking how many cells that had been in S-phase (EdU+) had transited the cell cycle and now had a 2N, G1 DNA content. At 6 hours after labeling, there was an ~50% reduction in EdU+ G1 cells in the USP37-depleted populations, suggesting that USP37-depleted cells proceed slowly through S-phase (Figure 1B). However, a requirement for USP37 in efficient progression through mitosis has also been reported, which could delay transition of the EdU+ cells to G1 in this context (Yeh et al., 2015). To confirm that delayed S-phase progression contributed to the altered cell cycle dynamics rather than mitotic defects, we performed dual labeling experiments in asynchronous U2OS cells to determine whether USP37-depleted cells indeed spend more time in S-phase. Cells were labeled with IdU to mark replicating cells. After an 8 hour chase, replicating cells were again identified by pulse labeling with CldU. The incorporation of CldU in the IdU+ population was analyzed to determine the fraction of cells that remained in S-phase. In keeping with a role for USP37 in S-phase progression, USP37-depleted cells demonstrated a modest, but reproducible, increase in the percentage of cells remaining in S-phase (IdU+,CldU+, Figures 1C-D).

**Figure 1.**
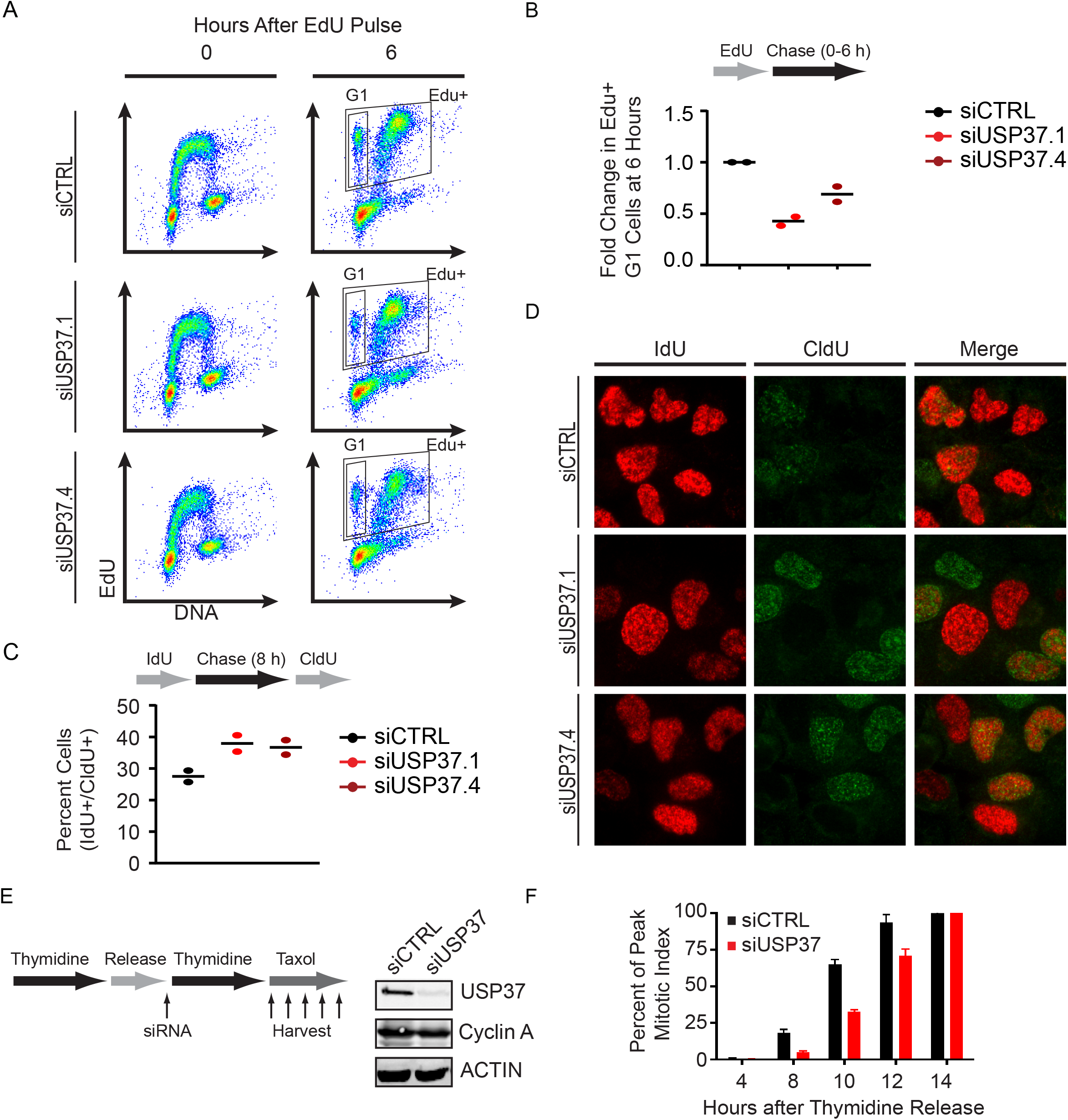
USP37 Promotes Progression through S-phase. (A) Schematic representation of the experimental scheme (top), quantification of the fold-change in EdU-positive (EdU+) HCT116 cells that had transited to G1 at 6 hours-post labeling (bottom). Individual data points for 2 experiments is shown. (B) Representative flow cytometry analyses for an experiment from (A). (C) Schematic representation of the experimental scheme (top), quantification of the percent of in IdU-positive (IdU+) U2OS cells that incorporated CldU (CldU+) 8 hours later (bottom). Individual data points for 2 experiments is shown. (D) Representative imgages for an experiment from (C). (E) Schematic representation of USP37-depletion/synchronization scheme, left. HeLa cells, treated as in E, were analyzed by immunoblot for the indicated proteins. (F) HeLa cells, treated as in E, were analyzed for phosphorylated Histone H3 Ser10, as a marker for mitosis, at the indicated timepoints.

Because Cyclin A accumulation was delayed, but not abolished in USP37-depleted cells, and CDT1 is not regulated by USP37 in S-phase, we reasoned that synchronizing these cells at G1/S would allow Cyclin A accumulation and synchronous S-phase entry (Hernandez-Perez et al., 2016; Huang et al., 2011). Given that USP37 is degraded during G2 and late mitosis/G1 by the SCF^βTRCP^ and APC/C^CDH1^ ligases, respectively, we transfected HeLa cells with control or USP37-targeting siRNAs just prior to the intervening mitosis of a double thymidine block to prevent the accumulation of USP37 during the subsequent G1 phase (Figure 1E, left panel). Indeed, Cyclin A accumulated to comparable levels in both populations by the end of the second block (Figure 1E, right panel). If USP37 acts only to promote the timely entry into S-phase then cell cycle progression should proceed similarly in control and USP37-depleted cells from this point. After release from thymidine we treated cells with taxol to trap cells in mitosis and monitored the appearance of the mitotic marker phospho-histone H3 S10 over time. USP37-depleted cells exhibited a marked delay in mitotic accumulation, suggesting that these cells were transiting slowly through S-phase (Figure 1F). Together our results suggest that replication kinetics are diminished in the absence of USP37.

### USP37 Loss Leads to Replication Stress

Our results thus far indicate that USP37 promotes efficient progression of the replication process. We noticed that the delays in progression through S-phase appeared to be more pronounced in the synchronized population. While synchronization via thymidine blocks is widely used to study cell cycle progression and generally considered innocuous, it does cause replication stress (Kurose et al., 2006). Thus our observation suggested that USP37-depleted cells may be undergoing and/or sensitive to replication stress. We thus examined the impact of USP37 loss on the emergence of DNA damage in replicating cells. Examination of γH2AX (Figures 2A-B) or 53BP1 (Figures 2E-F) foci in USP37-depleted cells did not indicate significant increases in replication stress/DNA damage in U2OS and Hela cells. However, upon treatment with low doses of the replication poison aphidicolin there is a significant increase in the number of foci for both of these markers in the absence of USP37 (Figures 2A, C, D, F). We next depleted USP37 from synchronized cells as in Figure 1E and analyzed replication stress/DNA damage as above. Consistent with previous reports, double-thymidine blocked cells exhibited increased γH2AX foci, which were similar between control and USP37-depleted cells (Figures 3A-B) (Kurose et al., 2006). Indeed, labeling cells with EdU demonstrated that USP37-depleted cells initiate replication upon release from the thymidine block comparably to control-depleted cells (Figure S1), consistent with the accumulation of Cyclin A in synchronized cells (as in Figure 1E). However, whereas the number of γH2AX foci per nucleus was reduced as cells began replication in control populations, the number of foci in USP37-depleted cells remained elevated. These analyses revealed that USP37-depleted cells exhibit increased levels of DNA damage in early S-phase upon release from thymidine. Similarly, double-thymidine blocked cells also exhibited elevated levels of 53BP1 foci. Upon release from the thymidine block the number of 53BP1 foci in USP37-depleted cells increased as cells underwent replication, in contrast to control cells (Figures 3C-D). Together these data suggest that USP37 promotes proper progression through S-phase and that the loss of USP37 leads to enhanced sensitivity to replication stress.

**Figure 2.**
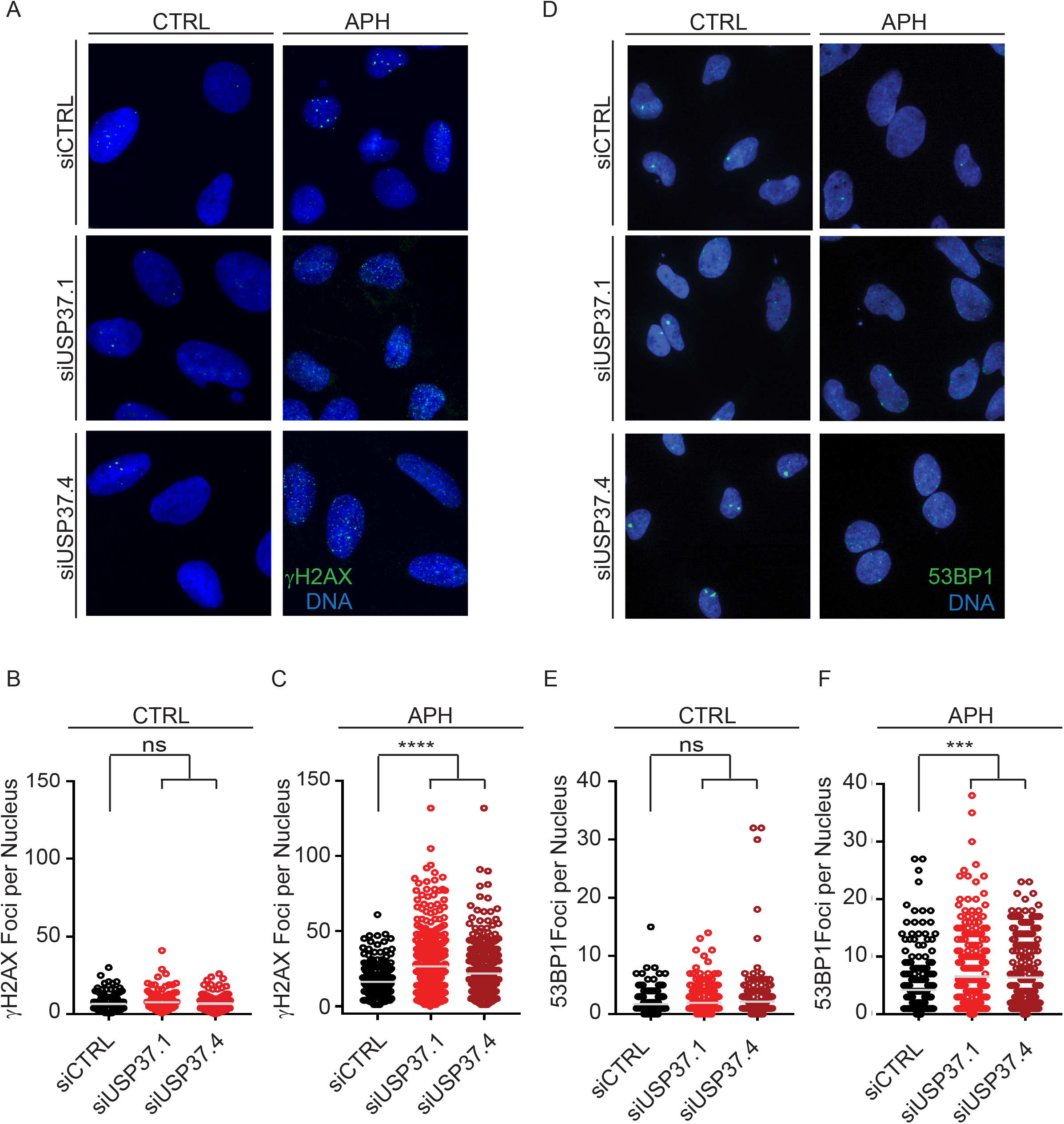
USP37-Depleted Cells Display Increased Sensitivity to Replication Stress. (A) U2OS transfected with the indicated siRNAs were analyzed for the formation of γH2AX foci. (B) Quantification of the number of foci per nucleus cells, as in A, under control conditions, n>140 cells per condition (C) Quantification of the number of foci per nucleus cells, as in A, after treatment with 200 nM aphidicolin (APH), n>270 cells per condition. The mean foci number is indicated. (D) HeLa transfected with the indicated siRNAs were analyzed for the formation of 53BP1 foci. (E) Quantification of the number of foci per nucleus cells, as in D, under control conditions, n>200 cells per condition (F) Quantification of the number of foci per nucleus cells, as in D, after treatment with 200 nM aphidicolin (APH), n>200 cells per condition. The mean foci number is indicated. For all experiments data were analyzed by one-way ANOVA with Dunnet’s post-test; ***,p<0.001, ****, p<0.0001

**Figure 3.**
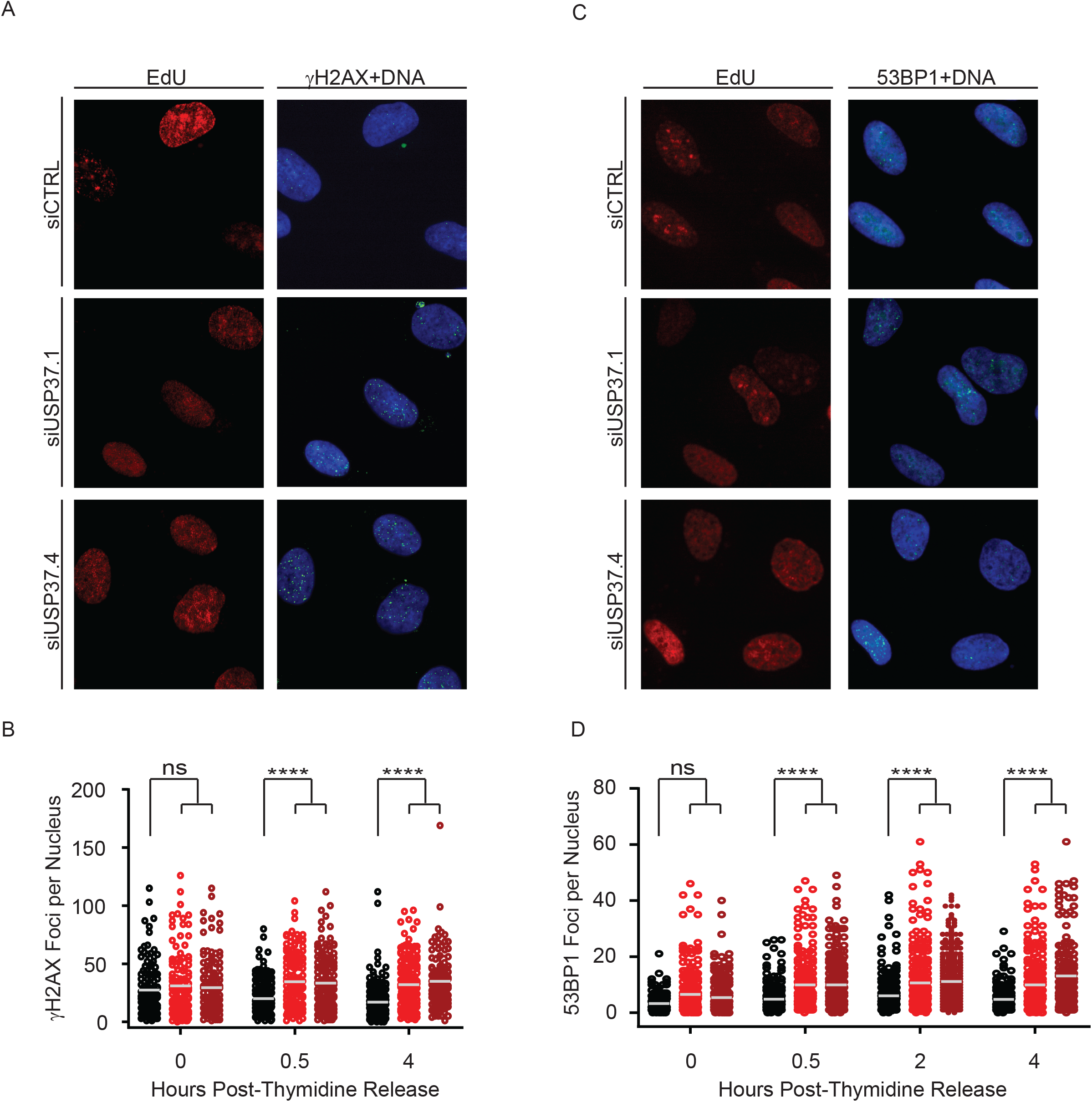
USP37-Depleted Cells Display DNA Increased Damage Markers During Perturbed Replication. (A) U2OS transfected with the indicated siRNAs as in (Figure 1A) were analyzed for the formation of γH2AX foci. Images from 4 hours post thymidine-release are shown. (B) Quantification of the number of foci per nucleus cells, as in A, n>120 cells per condition. The mean foci number is indicated. (C) HeLa transfected with the indicated siRNAs as in (Figure 1A) were analyzed for the formation of 53BP1 foci. Images from 4 hours post thymidine-release are shown. (D) Quantification of the number of foci per nucleus cells, as in D, n>150 cells per condition. The mean foci number is indicated. For all experiments data were analyzed by one-way ANOVA with Sidak’s post-test; ***,p<0.001, ****, p<0.0001

### USP37 Promotes Replication Stress Tolerance

To gain further insight into the biological impact of the increased sensitivity to replication stress upon depletion of USP37, we examined the proliferation of cells with and without replication stress. Real-time proliferation analysis were performed on USP37-depleted HCT116 and Hela cells with and without addition of aphidicolin to induce stress. In both cases, USP37-depleted cells proliferated more slowly than the control populations, although this effect was less pronounced in HeLa cells (Figures 4A-B). Moreover, in line with the increased levels of replication stress, USP37-depleted cells in both cell lines showed significant loss of proliferation upon induction of replication stress in comparison to both control cells undergoing stress and USP37-depleted cells in the absence of induced stress. Further analysis revealed a dose dependent response in HCT116 cells (Figures S2A-C). To extend these findings, we examined the effects of USP37 depletion in multiple cell lines. Similar sensitivity was observed in a third cell line, H1299, with hydroxyurea as the stress-inducing agent (Figure S2D). Two additional cell lines, MCF7 and MDA-MB-468, exhibited clearly diminished proliferation upon USP37-depletion in the absence of drug with MCF7 exhibiting marked sensitivity (Figures S2E-F). We then examined the impact of USP37-depletion on the long-term proliferative capacity of cells. Consistent with our kinetic growth analyses, depletion of USP37 caused a reduction in clonogenic growth in HCT116 (Figures 4C-D) and H1299 (Figures S3A-B) cells. The growth defect was significantly exacerbated by the induction of replication stress by hydroxyurea in comparison to both USP37-depleted cell proliferation in the absence of HU and control cells treated with HU. Similar results were also obtained in U2OS cells (Figures S3C-D). To more directly compare the impact of USP37-depletion on the sensitivity to replication stress, we treated control or siUSP37-transfected HCT116 and U2OS cells with thymidine, HU or APH to induce replication stress and examined colony forming potential in comparison to growth in the absence of treatment. USP37-depleted HCT116 cells displayed significant loss of proliferation upon induction of replication stress by all three agents, whereas control cells exhibited significant sensitivity only to APH, although to a lesser degree than the USP37-depleted cells (Figures 4E-F). U2OS cells behaved similarly. Control U2OS cells were modestly, but not significantly sensitive to replication stress induced by all three agents while USP37-depleted cells were significantly sensitive to both HU and APH. And, although it did not reach statistical significance, the colony formation capacity of USP37-depleted U2OS cells upon exposure to thymidine was dramatically diminished relative to control-depleted cells (Figures S3E-F).

**Figure 4.**
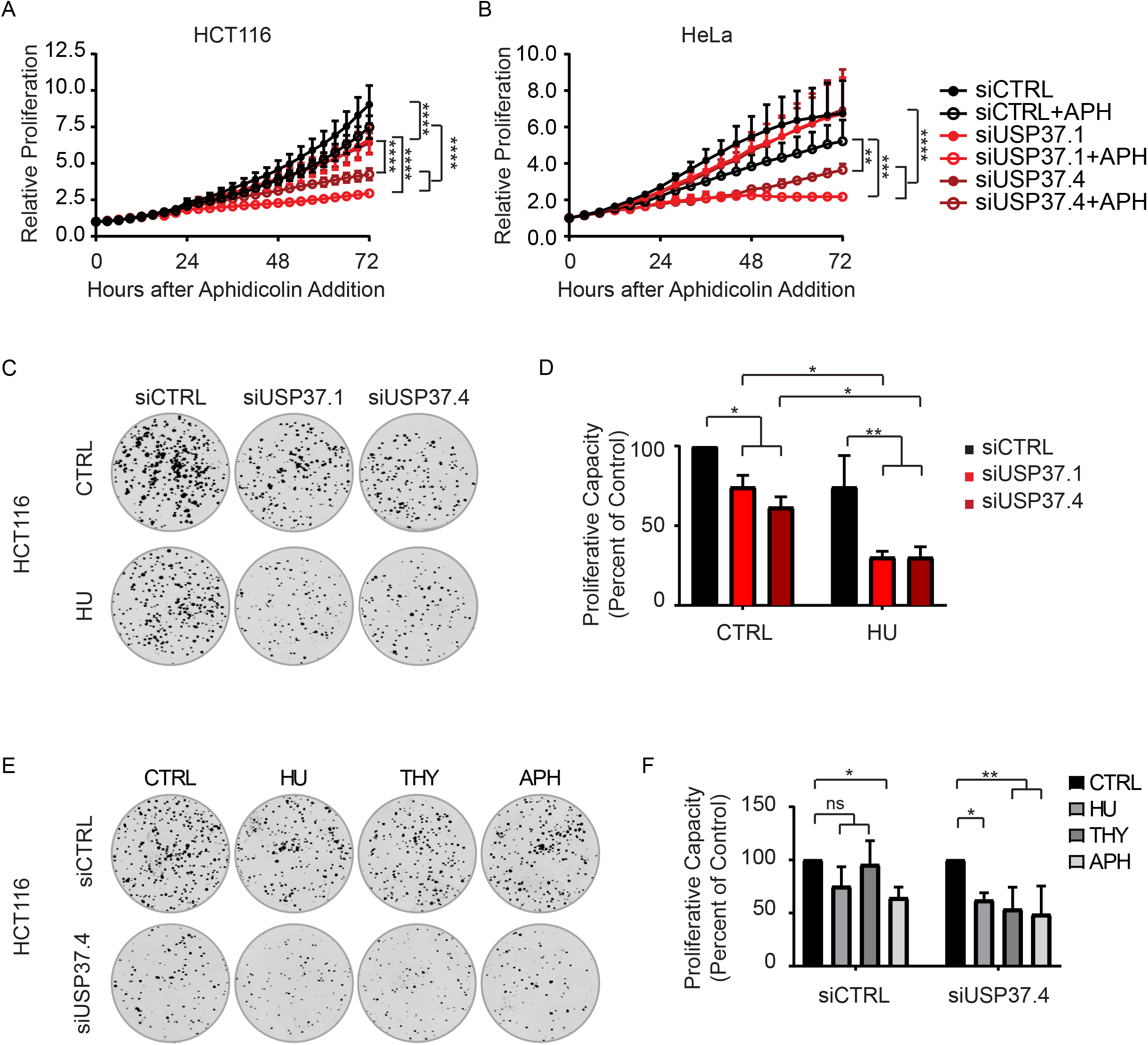
USP37 Promotes Proliferation and Tolerance of Replication Stress. (A) Cell proliferation (normalized to t=0) was analyzed in HCT116-H2BGFP cells transfected with the indicated siRNAs ± 100 nM APH. Data from two independent experiments, performed in triplicate. Mean and standard deviation are indicated. Data were analyzed by two-way ANOVA with Tukey’s post-test; ****, p<.0001. (B) Cell proliferation (normalized to t=0) was analyzed in HeLa cells transfected with the indicated siRNAs ± 200 nM APH. Data from two independent experiments, performed in triplicate. Mean and standard deviation are indicated. Data were analyzed by two-way ANOVA with Tukey’s post-test; **,p<0.01, ***,p<0.001, ****, p<0.0001. (C) Colony formation of HCT116 cells transfected with the indicated siRNAs ± treatment with 1 mM hydroxyurea (HU) for 18 hours. (D) Quantification of data as in (C), normalized to control, no drug. Data from two independent experiments, performed in triplicate. Mean and standard devation are indicated. Data were analyzed by two-way ANOVA with Tukey’s post-test; *,p<0.05, **,p<0.01, (E) Colony formation of HCT116 cells transfected with the indicated siRNAs ± treatment with 1 mM HU, 200 mM thymidine (THY), or 7 μM APH for 18 hours. (F) Quantification of data as in (E), normalized to control conditions, for each siRNA. Data from three independent experiments, performed in triplicate. Mean and standard deviation are indicated. Data were analyzed by two-way ANOVA with Dunnet’s post-test; *,p<0.05,**,p<0.01,

We next extended our analyses of the effect of USP37-depletion on growth beyond cell lines and depleted USP37 in zebrafish embryos using morpholino technology. A translation-blocking morpholino efficiently depleted USP37 protein in embryos (Figure S4A). In keeping with our cell line data, USP37-deficient embryos exhibited a survival deficit, along with increased cellular death that was exacerbated by replication stress induced by hydroxyurea (Figures S4B-D). Depletion of USP37 in embryos also resulted in developmental defects that manifested as a body curvature phenotype, which was significantly enhanced by replication stress (Figures S4C, E-F). Notably, increased body curvature has been associated with genomic instability and sensitivity to replication stress/DNA damage in developing zebrafish (Dimri et al., 2015; Ghiselli, 2006; Ishaq et al., 2013; Langheinrich et al., 2002; Robu et al., 2012). Together, our results indicate that USP37-deficient cells are generally sensitive to replication stress.

### UPS37 Promotes CHK1 Activity

Given that USP37-depleted cells exhibit slower progression through S-phase and delayed mitotic entry, it is possible that these cells exhibit a higher basal level of replication stress and thus replication checkpoint activity. However, the fact that we do not observe evidence of DNA damage in the absence of replication stress as well as increased sensitivity to replication stress, suggests that these cells may have defects in the checkpoint response. To gain insight into the status of the replication checkpoint in the absence of USP37 we examined phosphorylation of the essential replication checkpoint effector kinase CHK1 as Ser345 (pS345), which is indicative of its full activation and is frequently used to measure checkpoint activation (Wang et al., 2012). We first examined the kinetics of CHK1 activation by pulsing the cells with HU. We did not observe a heightened level of basal CHK1 phosphorylation in USP37-depleted cells (Figure 5A). However, although CHK1-pS345 appeared with similar kinetics and accumulated over time in both control and USP37-depleted cells, we noted that the overall levels of CHK1-p345 were lower in the absence of USP37 (Figures 5A-B). The lag in CHK1 phosphorylation could result from a defect in activation of the kinase. Interestingly, the CHK1 activating phosphorylation events elicit a conformational change that not only places the kinase domain into an active conformation, but also exposes degrons that prompt the destruction of active CHK1 to limit activation and promote efficient checkpoint recovery (Huh and Piwnica-Worms, 2013; Leung-Pineda et al., 2009; Zhang et al., 2009; Zhang et al., 2005). A similar reduction in pCHK1 levels was observed after prolonged replication stress that was accompanied by a 25% reduction in the level of total CHK1 (Figure 5C-D). Together these data indicate that USP37-depleted cells fail to maintain active CHK1 during replication stress.

**Figure 5.**
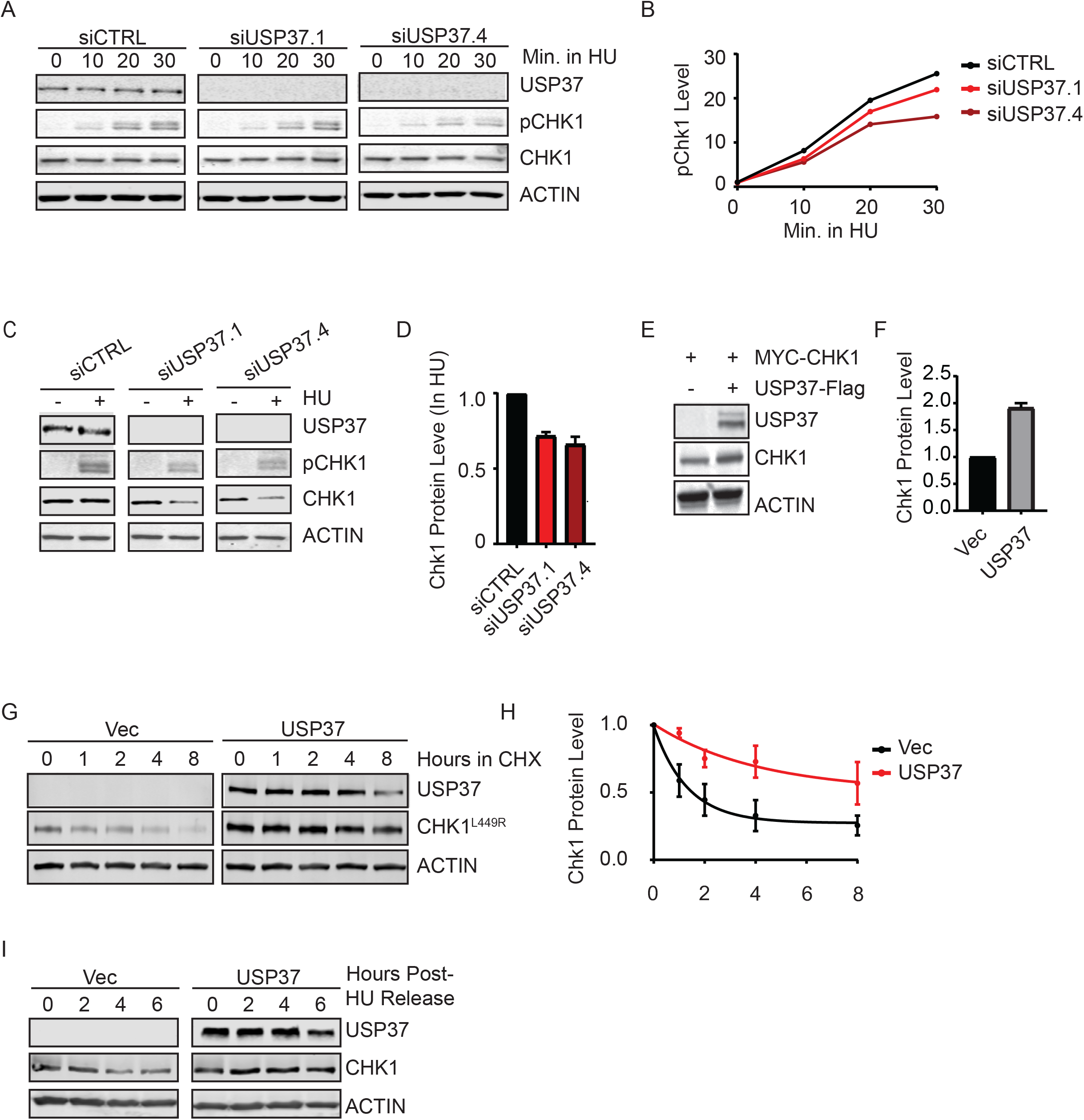
USP37 Promotes CHK1 Stability. (A) HCT116 cells were transfected with the indicated siRNAs and examined by immunblotting at the indicated times after treatment with 10 mM HU. (B) Quantification of CHK1 phosphorylation at S345 (pCHK1), normalized to total CHK1 levels. (C) HCT116 cells transfected with the indicated siRNAs were analyzed by immunoblot 18 hours after treatment with 2 mM HU. (D) Quantification of CHK1 protein levels after HU treatment. Data from two-independent experiments is shown. Mean and standard deviation are indicated. (E) 293T cells were transfected with plasmids encoding MYC-CHK1 ± FLAG-USP37 and analyzed by immunoblot. (F) Quantitation of data as in (E) showing the fold increase in MYC-CHK1 levels. Data is from three independent experiments. Mean and standard deviation are indicated. (G) 293T cells were transfected with plasmids encoding constitutively active MYC-CHK1^L449R^ ± FLAG-USP37 and analyzed by immunblot after treatment with 100 mg/ml cycloheximide (CHX). (H) Quantification of data as in (G). Data is from three independent experiments. Mean and standard deviation are indicated. (I) 293T Cells were transfected with empty vector of a plasmid encoding FLAG-USP37 and treated with 2mM HU for 16 hours. Protein levels were followed by immunoblot after release form HU.

To test the possibility that USP37 may affect CHK1 protein levels, we first asked whether co-expression of USP37 could bolster the expression of ectopic CHK1 in 293T cells. The presence of USP37 resulted in a 2-fold increase in the levels of co-expressed CHK1, relative to vector alone (Figures 5E-F). Given this impact on ectopic protein these results suggest that USP37 may act upon CHK1 protein rather than by impacting transcription or translation. Accordingly, we asked whether USP37 promotes the stability of CHK1 using cycloheximide to block translation and monitored protein half-life. To directly test the impact of USP37 on CHK1 stability, we took advantage of the fact that the CHK1^L449R^ mutant is in the constitutively open conformation and is predicted to be inherently unstable, and allows us to test the ability of USP37 to regulate the stability of active CHK1 without replication stress and irrespective of any potential defects in CHK1 activation (Han et al., 2016). Co-transfection experiments in 293T cells revealed that expression of USP37 dramatically stabilizes CHK1^L449R^, extending the half-life of the protein from ~ 1 hour to ~2.8 hours (Figures 5 G-H). Similarly, expression of USP37 in 293T cells prevented the loss of Chk1 protein levels following release from hydroxyurea (Figure 5I). Together these data suggest that CHK1 may be a substrate of USP37.

### USP37 Interacts with and Deubiquitinates CHK1

To further examine the possibility that CHK1 is a substrate of USP37 we sought to determine whether USP37 activity was required for its ability to impact CHK1 levels. Indeed, whereas expression of USP37 prevents the degradation of CHK1 following cycloheximide treatment, expression of the catalytically inactive USP37^C350A^ mutant failed to impact CHK1 levels (Figure 6A). Based on these observations, we asked whether USP37 and CHK1 interact. In an in vitro binding assay, recombinant GST-USP37 was able to interact with in vitro translated CHK1, but not the AURKB kinase, whereas GST-CDH1 was able to efficiently interact with its known substrate AURKB, indicating that the protein was correctly folded and suggesting that the USP37-CHK1 interaction was specific (Figure 6B). Surprisingly, CHK1 also interacted with GST-CDH1, which we subsequently determined to be a substrate of the kinase (Pal, et al manuscript submitted), further suggesting that the in vitro binding interaction was valid and specific. Similarly, immunoprecipitation of USP37 and CHK1 proteins after co-expression in 293T cells revealed that the proteins bind *in vivo* as well (Figures 6C-D). These results further indicated that CHK1 is a substrate of USP37, prompting us to perform *in vivo* deubiqutination analyses. We co-expressed MYC-CHK1 or MYC-AURKB with 6xHIS-Ub in 293T cells with or without USP37-FLAG and analyzed the ubiquitinated proteome, purified on Ni^2+^-beads, for the presence of the MYC epitope. Consistent with the results of our stability assays, expression of USP37 resulted in a nearly complete loss of CHK1-Ub conjugates (Figure 6E). The activity of USP37 toward CHK1 was specific as there were no impacts to bulk Ub-conjugates or to Ub-conjugates of the non-USP37 interacting kinase AURKB. Together these results identify CHK1 as a substrate of USP37.

**Figure 6.**
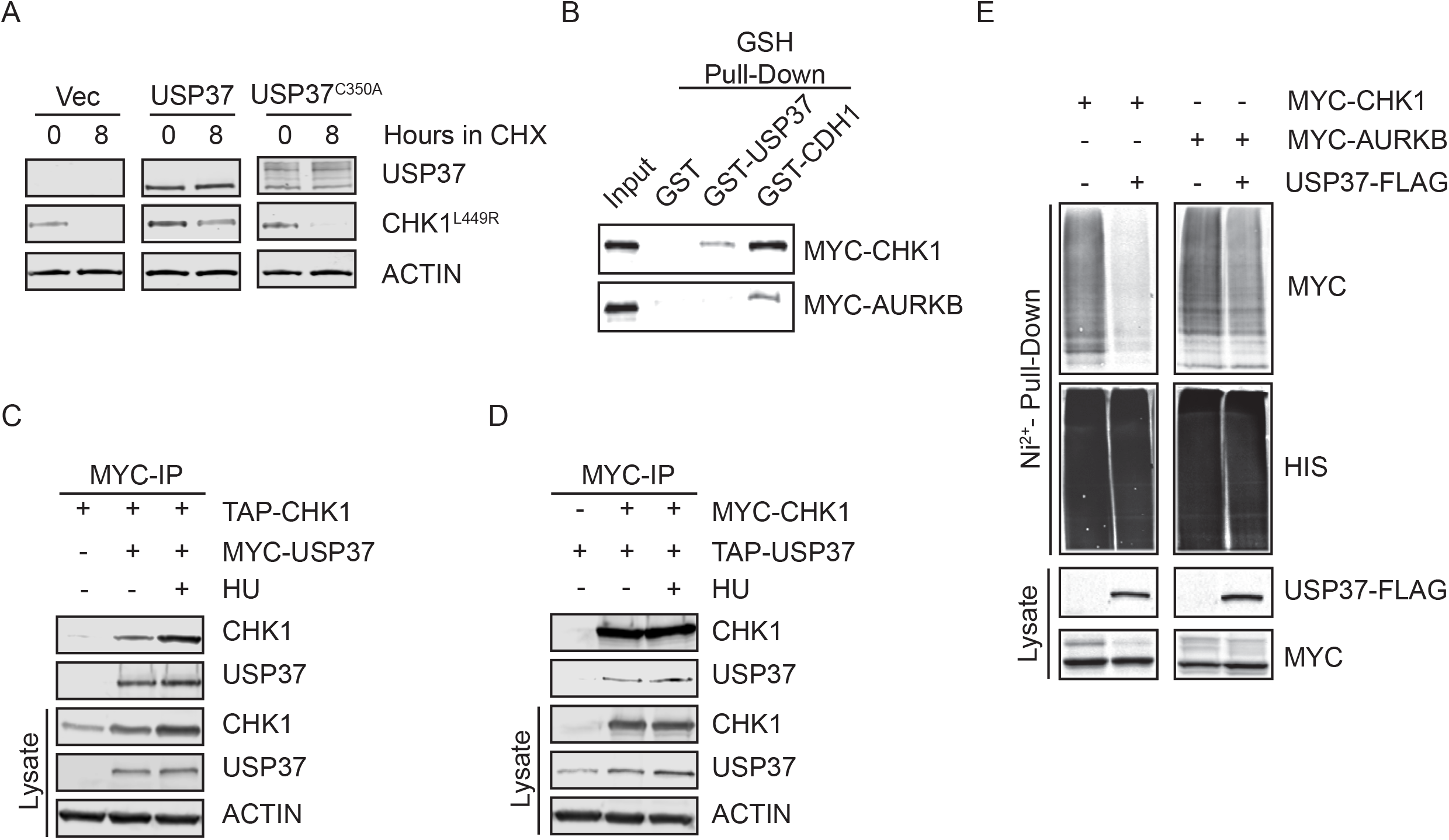
USP37 Binds and Deubiquitinates CHK1. (A) 293T cells were transfected with plasmids encoding MYC-CHK1^L449R^ ± FLAG-USP37 or the catalytically inactive USP37^C350A^ mutant and analyzed by immunoblot after treatment with 100 mg/ml cycloheximide (CHX). (B) Recombinant USP37 or CDH1 were utilized as bait in *in vitro* to capture *in vitro* translated MYC-CHK1 or MYC-AURKB. (C) 293T cells were transfected with plasmids encoding TAP-CHK1 ± MYC-USP37 as indicated and analyzed by immunoblot after MYC immunoprecipitation. (D) 293T cells were transfected with plasmids encoding MYC-CHK1 ± TAP-USP37 as indicated and analyzed by immunoblot after MYC immunoprecipitation. (E) 293T cells were transfected with plasmids encoding 6xHIS-Ub and MYC-CHK1 or MYC-AURKB ± FLAG-USP37. After treatment with MG132 to block the proteasome lysates and ubiquitin-conjugated proteins were analyzed by immunoblot.

## DISCUSSION

Here, we report the identification of CHK1 as a USP37 substrate. USP37 and CHK1 interact *in vitro* and *in vivo*. Expression of USP37 promotes steady state levels of CHK1, decreases CHK1-Ubiqutin conjugates, and stabilizes an active, unstable CHK1^L449R^ mutant. In the absence of USP37, CHK1 stability is compromised and cells mount an attenuated activation of CHK1. In keeping with the peak of USP37 expression and activity during late G1/S-phase, CHK1 joins the growing number of USP37 substrates (including Cyclin A, CDT1, and c-MYC) that are involved in the replication process and further implicates USP37 in the control of cell growth and the regulation of genome integrity (Figure 7) (Hernandez-Perez et al., 2016; Huang et al., 2011; Pan et al., 2015).

**Figure 7.**
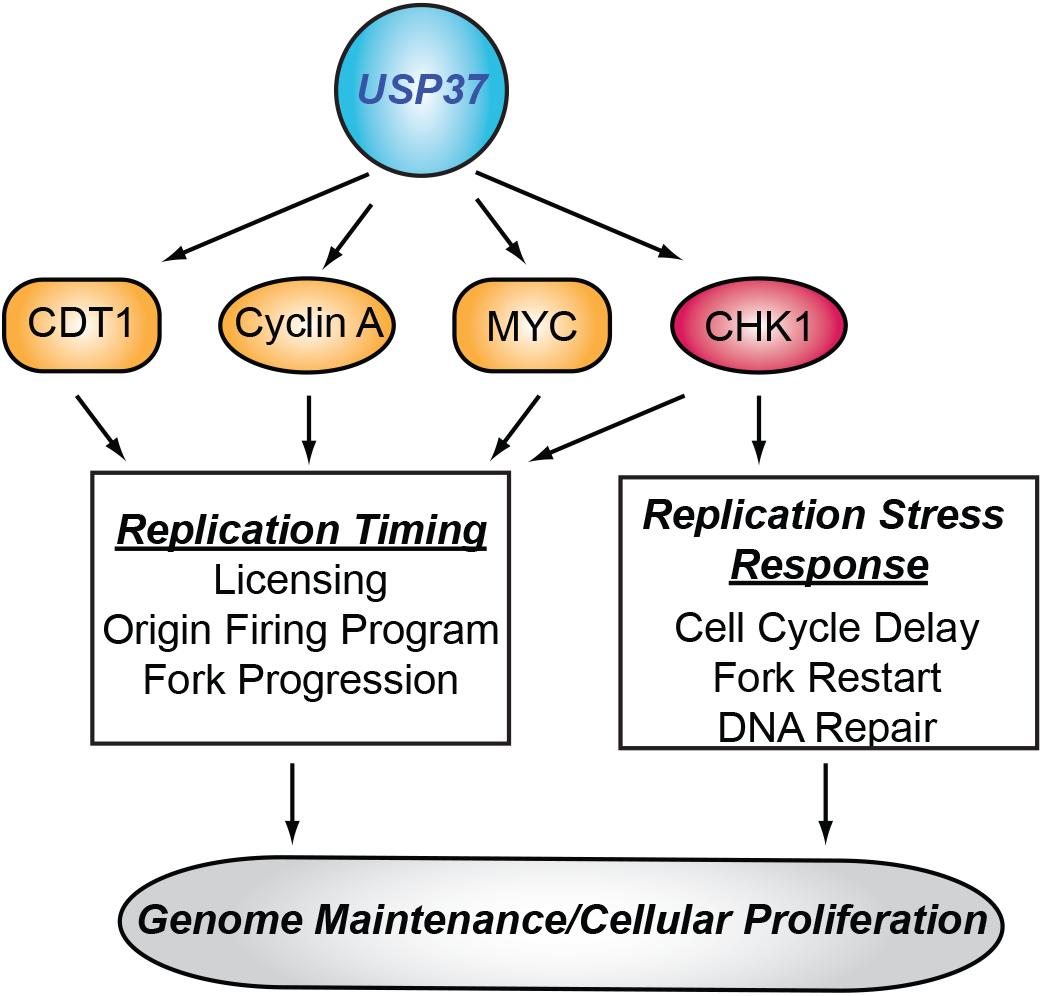
Current Model of the role of USP37 in the control of replication. USP37 is known to regulate the timing of S-phase by stabilizing factors involved in licensing (CDT1) and origin firing and fork progression (Cyclin A, MYC). By promoting CHK1 stability and activity, USP37 also strengthens the control of origin-firing and fork progression, while enhancing the response to perturbations in the replication process. See text for additional details.

The recent identification of CDT1 as a substrate of USP37 revealed that elevated USP37 could enhance MCM protein loading onto chromatin, consistent with its ability to promote S-phase (Hernandez-Perez et al., 2016). However, unexpectedly, depletion of USP37 was found to decrease replication fork speed and increase the number of replication origins fired. These observations are not anticipated for loss of CDT1, which would lead to fewer licensed origins and hence fewer origins capable of firing, a situation that permits increased fork speeds. In contrast, these observations are consistent with diminished CHK1 activity. CHK1 limits the number of forks that fire and the increased number of firing origins observed in the absence of CHK1 function causes a delay in replication fork speed as replication factors and nucleotides become limiting (Maya-Mendoza et al., 2007; Petermann et al., 2006; Petermann et al., 2010). Delayed fork speeds in CHK1 inhibited cells are not associated with delayed S-phase progression, in contrast to our observations (Petermann et al., 2010). However, the absence of CHK1 activity due to the use of inhibitors in these studies may mask the consequences of delayed progression. In contrast, slowing of replication fork rates has been associated with delayed cell cycle progression in other studies (Koundrioukoff et al., 2013; Lee et al., 2013). Notably, depletion of the replisome progression complex component AND-1, which also weakens CHK1 activity, attenuates fork speeds as well as delaying progression through S-phase (Hao et al., 2015; Yoshizawa-Sugata and Masai, 2009). Our data, together with this previous work suggest that regulation of CHK1 by USP37 promotes proper replication dynamics. However, we do not exclude the likely contribution of other yet to be identified USP37 substrates in the control of replication kinetics.

Alterations in replication dynamics are associated with replication stress. Thus it was somewhat unexpected that USP37-depleted cells did not exhibit signs of replication stress/DNA damage. However, similar alterations in replication kinetics and cell cycle progression without detection of classical markers of replication stress or the replication stress response have been reported (Koundrioukoff et al., 2013). Consistent with the idea that the USP37-depleted cells are undergoing replication stress these cells showed enhanced induction of γH2AX and 53BP1 foci after treatment with aphidicolin along with loss of proliferative capacity upon exposure to multiple forms of replication stress. These results are consistent with a recent study that found USP37-depleted cells are more sensitive to camptothecin (Qin et al., 2018). USP37 has been implicated in the regulation of DNA repair, primarily homologous recombination (HR) (Typas et al., 2015). Whereas defects in repair could contribute to the increased sensitivity to replication stress observed in the absence of USP37 and the observations with camptothecin, several observations suggest that increased susceptibility to insults to the replication process underlies the increased number of γH2AX and 53BP1 foci and subsequent failure to proliferate in these cells. First, USP37 has been found to interact with the replication machinery, indicating a role in the process, which is supported by altered replication timing and fork dynamics in USP37-depleted cells (Yeh et al., 2015). Second, the concentration of aphidicolin used in our live cell proliferation and foci analyses studies results in moderate fork slowing that is not associated with the formation of double strand breaks (Koundrioukoff et al., 2013). Third, upon release from a thymidine-induced replication block we observed an increase in 53BP1 foci in the absence of USP37 with γH2AX foci elevated similarly. Given that these foci were present in replicating cells, we interpret these foci to indicate stalled or malfunctioning forks. However, as the HR pathway is involved in the restart of stalled fork as well as the repair of collapsed forks, it is likely that USP37 deficiency leads to enhanced sensitivity by lowering the threshold for replication stress induction and impairing the mechanisms that allow for overcoming and recovering from challenges to the process. Future studies will determine the fate of replication forks and the nature of the lesions marked by γH2AX and 53BP1 in the absence of UPS37.

CHK1 activity is tightly regulated on multiple levels including the degradation of active CHK1 or of the upstream factor CLASPIN by the UPS (Bassermann et al., 2008; Faustrup et al., 2009; Gao et al., 2009; Huh and Piwnica-Worms, 2013; Leung-Pineda et al., 2009; Mailand et al., 2006; Mamely et al., 2006; Peschiaroli et al., 2006; Yuan et al., 2014; Zhang et al., 2009; Zhang et al., 2005; Zhu et al., 2014). Whereas multiple DUBs regulate CLASPIN, only two, USP1 and USP7, have been implicated in regulation of CHK1(Bassermann et al., 2008; Faustrup et al., 2009; Martin et al., 2015; McGarry et al., 2016; Yuan et al., 2014; Zhang et al., 2006; Zhu et al., 2014). Intriguingly, regulation of CHK1 by these DUBs may be context specific. USP1 indirectly regulates CHK1 by limiting ubiquitination of FANCD2/I and/or PCNA, which stimulates DDB1-dependent degradation of CHK1 (Guervilly et al., 2011). This mechanism may largely be important for types of damage that are dependent upon the Fanconi Anemia pathway for repair, including mitomycin C. Similarly, USP7 regulates CHK1 most efficiently under conditions that activate ATM, which results in the stabilization of ZEB1, a factor that enhances the USP7-CHK1 interaction (Alonso-de Vega et al., 2014; Zhang et al., 2014). Given that UPS37 expression and activity peak during S-phase, our findings suggest that USP37 regulates CHK1 during replication and, in particular, under conditions of replication stress. Increased replication stress, driven by oncogene activation, is inherent to cancer cells leading to an increased dependence on CHK1 and subsequently increased expression in cancers. Similar to CHK1, USP37 expression is elevated in multiple cancers. It will be interesting to determine whether the increased expression of USP37 in cancer reflects both an ability to drive replication as well as enhancing the ability to deal with replication stress caused by rapid cell cycle progression.

## EXPERIMENTAL MODEL AND SUBJECT DETAILS

### Cell Lines

293T, HCT116, HeLa, MDA-MB-231, MDA-MB-468, MCF7, T98G, and U2OS cells were all obtained from the American Type Culture Collection (ATCC). Cell lines are verified using the OSUCCC Genomics Shared Resource.

## REAGENTS and MATERIALS

(see supplemental table)

## METHOD DETAILS

### Cell Culture

Cell lines were obtained from the American Type Culture Collection. 293T, H1299, HCT116, HeLa, MDA-MB-231, MDA-MB-468, MCF7, T98G, and U2OS cells were cultured in Dulbecco’s modified Eagle’s medium (DMEM; Corning) supplemented with 10% FBS (Seradigm) and 1% Pen/Strep (GIBCO). All cells were incubated at 37°C and 5% CO_2_. T98G cells were synchronized by incubation in DMEM without FBS for 72 h and stimulated to re-enter the cell cycle by the addition of 20% FBS. For synchronization via double thymidine block, cells were treated with 2mM thymidine for 18 hours and released into fresh media after washing with PBS. After 9 hours, cells were again treated with 2mM thymidine for 18 hours and then released into S-phase by washing with PBS.

### Transfections and Treatments

siRNA transfections were performed using Lipofectamine RNAiMAX (ThermoFisher) according to the manufacturer’s protocol. Sequence of siRNAs used in this study are described in (). Plasmid transfections were performed using Mirus TransIT-LT1 reagent (Mirus Bio) according to the manufacturer’s protocol. Cycloheximide (Thermofisher) was used at a concentration of 100μg/mL for the indicated times. Aphidicolin, hydroxyurea and thymidine concentration and treatment duration are described in the figure legends.

### Molecular Cloning

cDNA encoding AURKB was cloned into pCS2+ MYC-DEST using Gateway Technology (Thermofisher).

### Immunoblotting

Cell extracts were generated in EBC buffer (50 mM Tris (pH8.0), 120mM NaCl, 0.5% Nonidet P-40, 1mM DTT and protease and phosphatase inhibitor tablets (ThermoFisher). Protein concentration was quantified by Pierce BCA assay (ThermoFisher) and samples were prepared by boiling in Laemmli buffer for 5 minutes. Equal amounts of whole cell lysates were resolved by hand-cast SDS-PAGE, transferred to PVDF membranes (Millipore). All blocking and primary antibody steps were performed in 5% nonfat dried milk diluted in TBST (137mM NaCl, 2.7mM KCl, 25mM Tris pH 7.4, 1% Tween-20), except for phospho-CHK1, which as incubated in 5% BSA diluted in TBST. All primary antibody incubations were performed with shaking at room temperature (for epitope tagged proteins) or at 4°C for 16 hours (for endogenous proteins). All secondary antibody incubations were performed with shaking at room temperature for 30 minutes in TBST + 0.02% SDS. Washing steps were performed using TBST and protein bands were visualized using the LI-COR Odyssey CLx infrared imaging system.

### Cellular Fractionation

Cell pellets were swelled on ice for 20 minutes in buffer A (10 mM HEPES pH 7.9, 10 mM KCl, 1.5 mM MgCl_2_) and protease inhibitors. After dounce homogenization, cells were pelleted at 1000xg for 3 min. The soluble fraction was collected and clarified at 20,000xg for 15 min. The pellet was washed with buffer B (3 mM EDTA, 0.2 mM EGTA, 1 mM DTT) and protease inhibitors for 30 minutes to remove nucleoplasmic protein and spun at 1000xg for 3 min. The pellet was extracted in EBC buffer with 500 mM NaCl to produce the chromatin bound fraction.

### Immunofluorescence, microscopy, and flow cytometry

Detection of DNA synthesis in proliferating cells was determined based on the incorporation of 5-ethynyl-2′-deoxyuridine (Click-IT EdU; Thermo Fisher Scientific) and its subsequent detection by a fluorescent azide through “click” chemistry per the manufacturer’s instructions. In brief, cells were pulsed for 15 minutes with 10 μM EdU (Thermo Scientific and fixed in 3.7% formaldehyde, and washed in PBS prior to EdU labeling by click chemistry. For detection of DNA damage U2OS or HeLa cells were seeded on glass coverslips and transfected with the indicated siRNAs. After 24 hours cells were treated overnight with vehicle or 200nM aphidicolin. Cells were fixed and permeablized with 0.5% Triton X-100 in PBS, washed and then blocked for 30 minutes at room temperature with 5% BSA in PBS. Cells were incubated with antibodies (1:500) in 5% BSA in PBST for 1 hour at room temperature. After washing the cells were incubated with Alexafluor secondary antibodies (1:500) in 5% BSA in PBST for 30 minutes at room temperature. DNA was counterstained with 1 μg/mL Hoechst 33342 and mounted with Fluoromount G (ThermoFisher). Cells were imaged using a Leica DM5500B fluorescent microscope as described previously. Images were analyzed and foci quantified with using FIJI. For combined EdU and DNA damage analysis, EdU detection was performed first as per the manufacturer’s instructions.

For flow cytometry, cells were incubated with the indicated siRNAs and harvested at the indicated times. For cell cycle progression studies, the cells were pulsed with EdU, as above, 48 hours after transfection. The cells were then harvested at the indicated times. Cells were fixed in ice-cold 70% ethanol and permeabilized in 0.1% Triton X-100 for 15 minutes on ice. EdU was visualized as above and the cells were incubated on with 200μg/ml RNAse A and 50μg/ml propidium iodide for 30min at 37 °C. Cells were then analyzed on a Becton Dickinson FacsCalibur instrument and analyzed with FlowJo software.

### Immunoprecipitations

Cells were transfected with the indicated plasmids and lysed in EBC buffer 48 hours after transfection. 1μg of the indicated antibody was mixed equal volumes lysate at 4°C overnight. Magnetic Protein A and Protein G beads (ThermoFisher) were then added and incubated at at 4°C for an additional 30 minutes. Beads were washed three times with EBC buffer and proteins were eluted by boiling beads in Laemmli buffer for 5 minutes and visualized by IB.

### *In Vitro* Binding Assay

pGEX6P1-USP37 or pGEX6P1-CDH1 were transformed into *Escherichia coli* BL21(DE3) cell. For expression of UPS37 bacteria were inoculated into a 20ml culture in Overnight Express media and grown for 8 hours at 37°C. The culture was used to inoculate two 1L cultures and grown for 15 hours at 20°C the OD_600_ was checked and the cells were grown for an additional 4 hours. OD_600_ was stable and the cells were incubated for an additional 8 hours and harvested. For expression of GST-CDH1, bacteria were grown in standard LB media to an OD_600_ of 0.5, and induced to express by addition of 0.1mM IPTG and incubated with shaking at 18 °C for 16 hours. Bacterial pellets were frozen in liquid nitrogen. Thawed cells were resuspended in 60mL lysis buffer (50mM Tris pH 8, 500mM NaCl, 0.5% Triton X-100, protease inhibitor tablets). GST-fusions were purified from crude lysate on glutathione sepharose (GE Lifesciences) following manufacturer’s protocol. Following elution, proteins were buffer exchanged (20mM HEPES pH 7.7, 100mM KCl, 1mM DTT and 5% glycerol) using Slide-a-lyzer dialysis units (ThermoFisher) following manufacturer’s protocols. MYC-CHK1 and MYC-AURKB were generated in a rabbit reticulocyte coupled transcription/translation system (Promega) following the vendor’s protocol.

For binding assays, 2mg of GST, GST-CDH1 or GST-USP37 were mixed with 10μl of IVT proteins on ice for 1 hour. Then, the proteins were diluted with TBST and mixed glutathione beads for 1 hour at 4 °C. Following four washes with TBST, protein complexes were eluted in Laemmli buffer and assessed by IB.

### S-phase Progression Assays

U2OS cells were seeded on glass coverslips and transfected with the indicated siRNAs. After 24 hours the cells were pulse labeled with 10μM IdU for 30 minutes and placed into fresh media. At 8 hours post-IdU, the cells were pulse-labeled with 10 μM CldU for 30min. THe cells were then fixed in 70% ethanol for 10 minutes at room temperature. After permeabilization in methanol chromatin was denatured with 1.5N HCl for 30 minutes, followed by blocking with BSA in PBST (PBS + 0.01% Tween-20). The coverslips were then incubated with sequentially with rat anti-BrdU, anti-rat secondary, mouse anti-BrdU, and anti-mouse secondary antibodies. All antibody incubations were followed by 3 washes in PBST. The DNA was counterstained with Hoecsht 33342 and coverslips were mounted with Flouromount G.

### Proliferation Assays

HCT116 H2B-GFP HeLa, H1299, MCF7, and MDA-MB-468 cells were seeded in 96 well plates. The cells were then transfected with the indicated siRNAs and incubated for 6 hours before the addition of drug, where indicated. Growth of triplicate samples was monitored by phase-contrast imaging or GFP count (HCT116 H2B-GFP cells) using the IncuCYTE Zoom (Essen Bioscience) system. Cells were imaged every 4 hours for 72 hours and growth was determined as fold increase of GFP+ cells or percent phase confluence at the start of imaging.

### Colony Formation Assay

HCT116, HeLa, H1299, MCF7 or U2OS cells were transfected with the indicated siRNA, then 24 hours later, seeded in triplicate for each condition in 6 well dishes. After 24 hours cells were treated with 1mM hydroxyurea, 200mM thymidine, or 2.5μg/ml aphidicolin for 18 hours. The cells were washed and given fresh media. Colonies were fixed and stained in 0.5% w/v crystal violet and 25% methanol after 10-14 days.

### *In Vivo* Deubiquitination Assay

293T cells were transfected with the indicated plasmids. After 24 hours the cells were treated with 10mM hydroxyurea and 100μM MG132 for 4 hours and then harvested. 90% of the cell suspension was lysed in (6M guanidine-HCL, 100mM Na_2_HP0_4−_ NaH_2_P0_4_, 10mM Tris-HCl pH 8.0, 5mM imidazole, 10mM β-mercaptoethanol) and 6xHIS-ubiquitinated proteins were captured on Ni-NTA resin (ThermoFisher) for 4 hours at room temperature. The beads were washed sequentially in lysis buffer (without imidazole), Buffer A pH 8 (8M urea, 100mM Na_2_HP0_4−_ NaH_2_P0_4_, 10mM Tris-HCl pH 8.0, 10mM β-mercaptoethanol), Buffer A pH 6.3 + 0.2% Triton X-100, Buffer A pH 6.3 + 0.1% Triton X-100 and eluted by boiling in Laemmli buffer. 10% of sample was used to prepare inputs. Pull-down eluates and inputs were separated on SDS-PAGE gels and analyzed by immunoblot.

### Protein Quantification and Stability

Protein bands were measured with Image Studio (LI-COR Bioscience), using median background subtraction. For determination of phosphor-CHK1 levels, signal intensity was normalized to the signal of the unmodified CHK1 in the same lane. For total CHK1 levels, signal intensity was normalized to the signal of ACTIN in the same lane. To analyze CHK1 stability, the protein level at time 0 was set to 1 and the fraction remaining was determined for subsequent timepoints. The protein half-life was determined using GraphPad Prism 7 to perform best-fit non-linear regression.

### Zebrafish Studies

Fish care and animal work were carried out in compliance with NIH guidelines for animal care and approved by the National Cancer Institute at Frederick Animal Care and Use committee (Study Proposal 17-416). Zebrafish used in these studies were wildtype AB.

Knockdowns for USP37 were carried out using the following translation blocking morpholino obtained from GENETOOLS, Philomath: USP37 5′-CCATCAGGGCGAAGATCCTCCACAA -3′. Embryos were injected with 250 μM of translation blocking morpholino at the one-cell stage using a microinjector PLI-90 (Harvard Apparatus). For hydroxyurea treatments, embryos were treated with 125 mM at 24hpf for 16h and examined at 72hpf. For acridine orange (AO) staining, embryos were placed at 24hpf in E3 medium with PTU as a pigment clearing method in presence or absence of 125 mM hydroxyurea. At 72 hpf, embryos were incubated for 1h in Acridine Orange (5 μg/ml) followed by 3 washes in E3 medium and imaged live in anesthetic reagent (MS-222) using a 4X water immersion objective (0.13 N.A) and a Nikon Eclipse Ni-E upright fluorescence microscope equipped with a DS-Ri2 camera.

For expression analysis, embryos were dechorionated and homogenized in lysis buffer (20mM Tris at pH 8, 137mM NaCl, 10% glycerol, 1% Triton X-100) with protease inhibitor cocktail (Roche), centrifuged for 10 min at 13,000 r.p.m and supernatants were boiled in sample buffer.

### Statistical Analysis

Statistical analyses were performed with GraphPad Prism Software using an unpaired Student’s T test 1-or 2-way ANOVA with Tukey or Dunnet’s post-test, respectively where appropriate (GraphPad Software, Inc.).

## Supporting information

Reagents and Materials

## ACKNOWLEDGEMENTS

This work was supported by The Ohio State University Comprehensive Cancer Center/Department of Radiation Oncology start-up funds and NIH grants R01 GM112895 and R01 GM108743 to MKS. Research reported in the publication was supported by The Ohio State University Comprehensive Cancer Center and the National Institutes of Health under grant number P30 CA016058. The authors thank The Ohio State University Comprehensive Cancer Center’s Genomics Shared Resource for technical support. We also thank Monica Venere and members of the Summers laboratory for insightful discussion and constructive comments on the manuscript. The content of this work is solely the responsibility of the authors and does not necessarily represent the official views of the National Institutes of Health.

## AUTHOR CONTRIBUTIONS

M.S., A.C.B., A.D.T., designed, performed and analyzed most experiments. B.S. performed some of the proliferation assays and *in vivo* interaction studies. D.P. A.D. performed the *in vitro* binding studies. D.P. also performed cellular assays. C.I. and C.J.W designed, analyzed zebrafish studies, which were performed by C.I. M.K.S. designed and performed experiments, helped in data analysis, and wrote the manuscript. All authors reviewed and approved the manuscript.

---

**Figure S1. USP37-Depleted Cells Enter S-Phase with Control-Depleted Cells After Release From Thymidine, Related to Figure 3.**

HeLa cells, treated as in Figure 1E were pulsed with EdU 30 minutes after release from the second thymidine block. The number of cells in S-phase (EdU+) in each population are indicated.

**Figure S2. USP37 Promotes Short-term Proliferation and Tolerance of Replication Stress, Related to Figure 4**

(A) Cell proliferation (normalized to t=0) was analyzed in HCT116-H2BGFP cells transfected with the indicated non-targeting siRNA (siCTRL) ± the indicated concentrations of APH. Data from two independent experiments, performed in triplicate. Mean and standard deviation are indicated.

(B) Cell proliferation (normalized to t=0) was analyzed in HCT116-H2BGFP cells transfected with the indicated USP37-targeting siRNA (siUSP37.1) ± the indicated concentrations of APH. Data from two independent experiments, performed in triplicate. Mean and standard deviation are indicated.

(C) Cell proliferation (normalized to t=0) was analyzed in HCT116-H2BGFP cells transfected with the indicated USP37-targeting siRNA (siUSP37.4) ± the indicated concentrations of APH. Data from two independent experiments, performed in triplicate. Mean and standard deviation are indicated.

(D) Cell proliferation (normalized to t=0) was analyzed in H1299 cells transfected with the indicated siRNAs ± 100 μM HU. Data is from an experiment, performed in triplicate. Mean and standard deviation are indicated.

(E) Cell proliferation (normalized to t=0) was analyzed in MDA-MB-468 cells transfected with the indicated siRNAs. Data is from an experiment, performed in triplicate. Mean and standard deviation are indicated.

(F) Cell proliferation (normalized to t=0) was analyzed in MCF7 cells transfected with the indicated siRNAs. Data is from an experiment, performed in triplicate. Mean and standard deviation are indicated.

**Figure S3. USP37 Promotes Colony Formation and Tolerance of Replication Stress, Related to Figure 4**

(A) Colony formation of H1299 cells transfected with the indicated siRNAs ± treatment with 1 mM HU for 18 hours.

(B) Quantification of data as in (A), normalized to control, no drug. Data from three independent experiments, performed in triplicate. Mean and standard deviation are indicated. Data were analyzed by two-way ANOVA with Tukey’s post-test; *,p<0.05, **,p<0.01, ***, p<0.001

(C) Colony formation of U2OS cells transfected with the indicated siRNAs ± treatment with 1 mM HU for 18 hours.

(D) Quantification of data as in (C), normalized to control, no drug. Data from an experiment, performed in triplicate. Mean and standard deviation are indicated.

(E) Colony formation of U2OS cells transfected with the indicated siRNAs ± treatment with 1 mM HU, 200 mM, THY, or 7 μM APH for 18 hours.

(F) Quantification of data as in (F), normalized to control conditions for each siRNA. Data from three independent experiments, performed in triplicate. Mean and standard deviation are indicated. Data were analyzed by two-way ANOVA with Dunnet’s post-test; *,p<0.05.

**Figure S4. USP37 Promotes Growth Tolerance of Replication Stress in Zebrafish Embryos, Related to Figure 4**

(A) USP37 protein expression was examined by immunoblot after injection with a USP37 translation-blocking morpholino (TB MO).

(B) Analysis of death in 72 hpf embryos uninjected or injected with USP37 TB MO and untreated or treated with 125 mM hydroxyurea for 16 hours at 24hpf. These data are from 2 independent experiments with total numbers of fish counted as follows; Uninj. −HU, n=71; Uninj. +HU, n=90; TB −HU, n=47; TB +HU, n=51. One-way ANOVA was applied to obtain p values. ** p<0.01

(C) Fish, as in (B), treated with 125 mM HU for 24 hours and stained with acridine orange to visualize apoptotic cells. Arrowheads indicate the regions of increased apoptosis visualized in the tail.

(D) Representative images of tails of fish from two different groups (upper and lower panels), examined as in (C), from 3 independent experiments, as in (C).

(E) Representative brightfield images of live 72 hpf embryos uninjected or injected with USP37 TB MO ± HU.

(F) Quantification of body curvature defects in embryos as in (E). Mean and SD from 4 independent experiments with total numbers of fish counted as follows; Uninj. −HU, n=62; Uninj. +HU, n=73; TB −HU, n=38; TB +HU, n=28. One-way ANOVA was applied to obtain p values. ** p< 0.005, **** p<0.0001

