## Supplementary figures and images for "The Deubiquitinating Enzyme USP37 Stabilizes CHK1 to Promote the Cellular Response to Replication Stress"

### Supplemental Figures

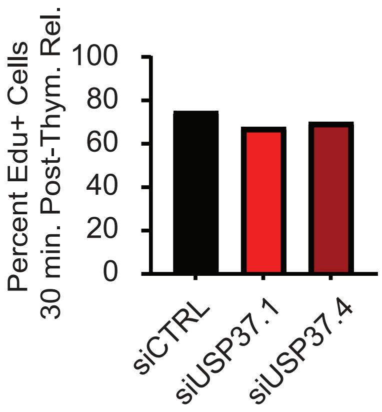

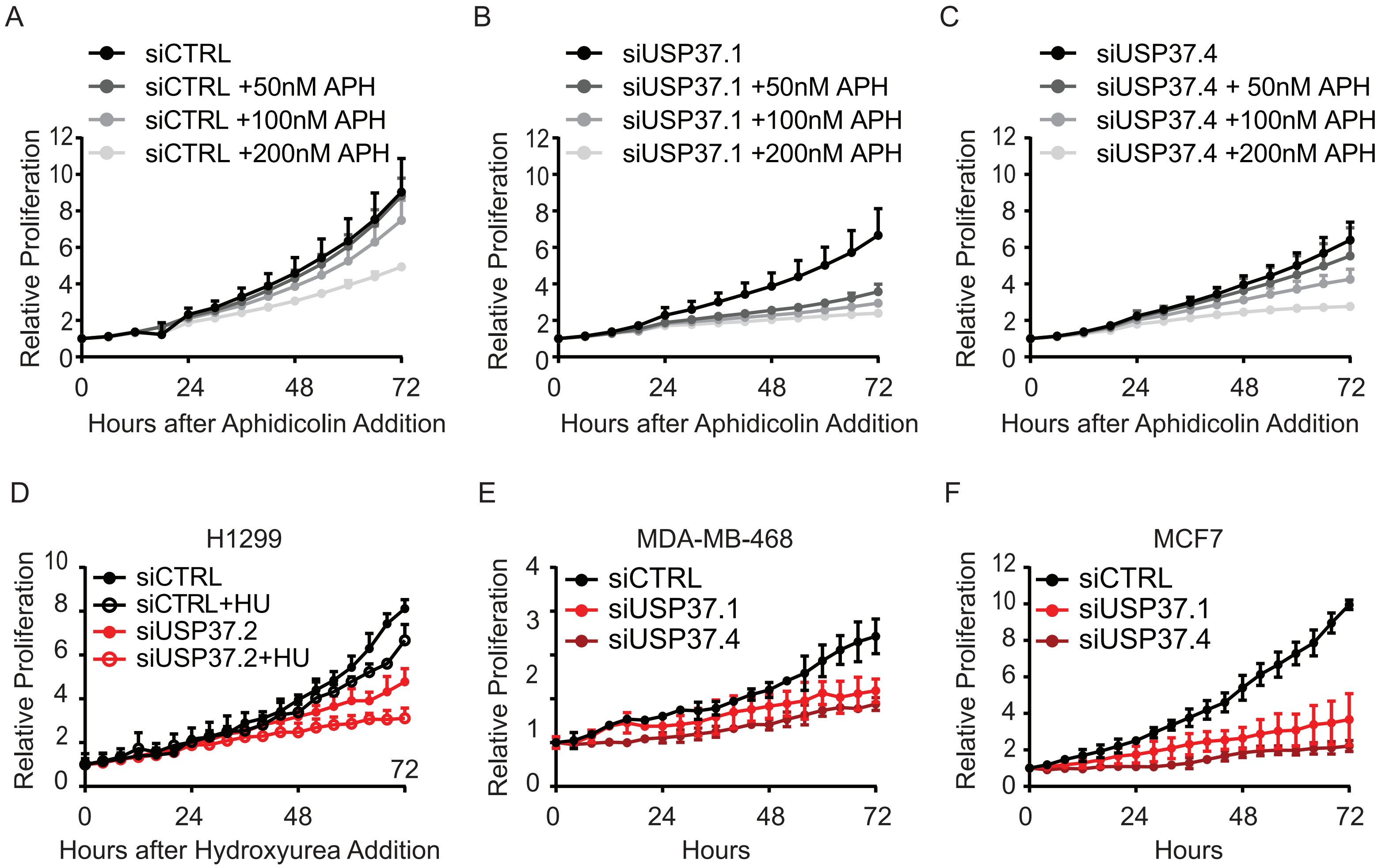

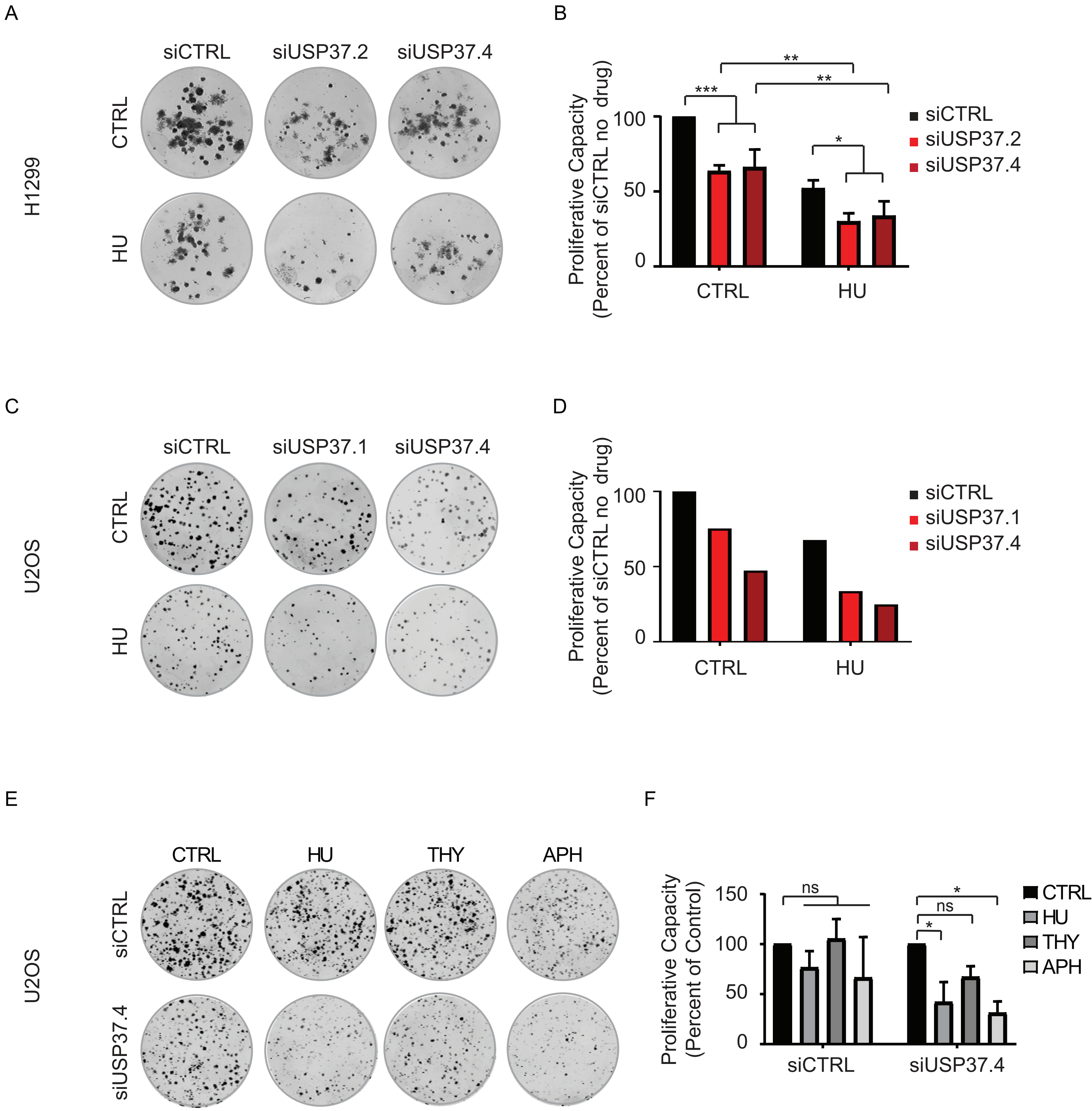

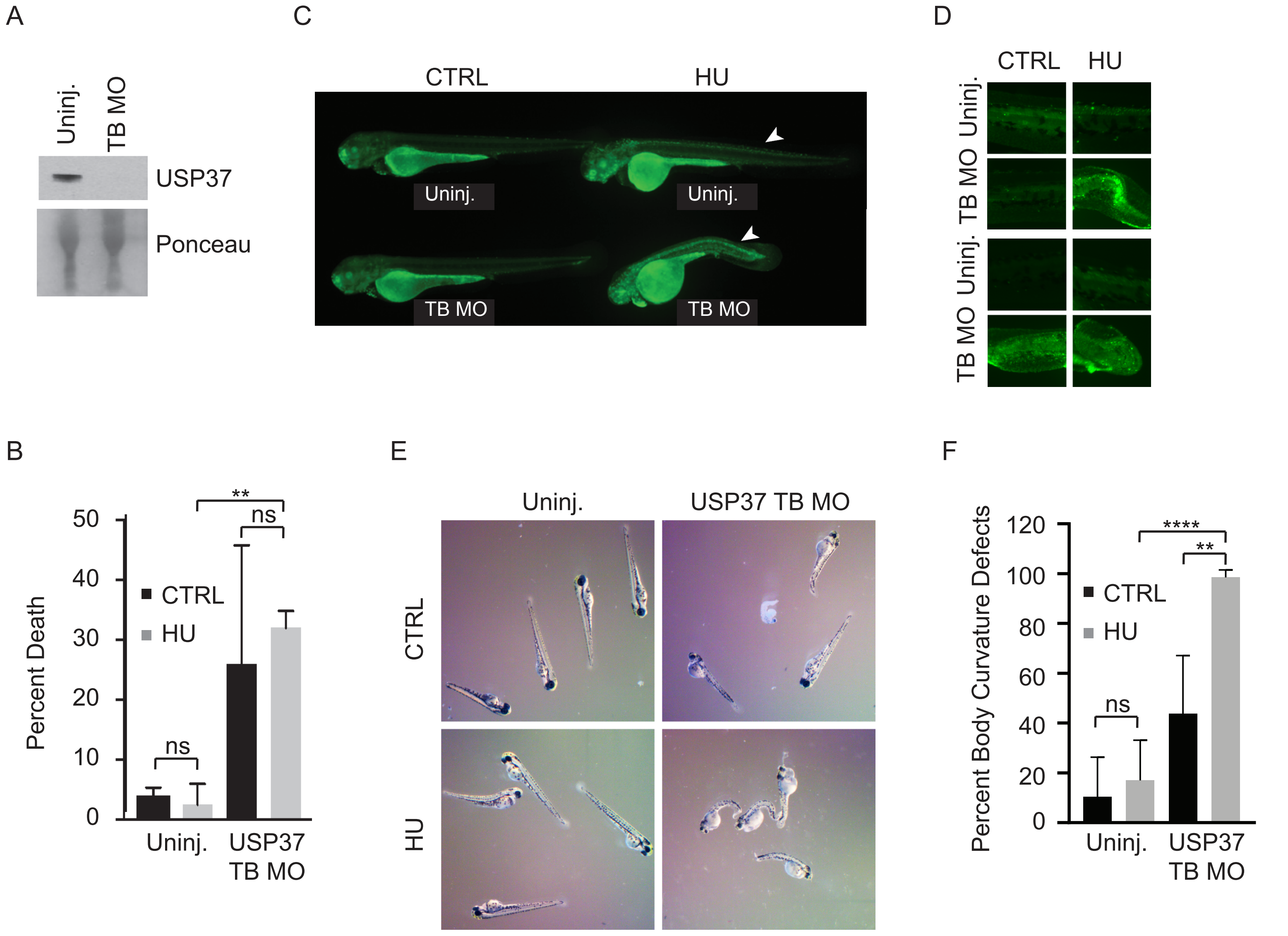
