## Supplementary material for "The Deubiquitinating Enzyme USP37 Stabilizes CHK1 to Promote the Cellular Response to Replication Stress": Reagents and Materials

| REAGENT or RESOURCE | SOURCE | IDENTIFIER |
|---|---|---|
| **Antibodies** | | |
| 53BP1 | BD Biosciences | Cat# 612522, RRID:AB_2206766 |
| BrdU | ThermoFisher | Cat# MA1-82088, RRID:AB_927214 |
| BrdU | BD Biosciences | Cat# 347580, RRID:AB_400327 |
| Chk1 | CST | |
| CHK1 | Santa Cruz | Cat# sc-8408; RRID:AB_627257 |
| CHK1 phospho-S345 | CST | Cat# 2348S; RRID:AB_331212 |
| Cyclin A1 | Santa Cruz | Cat# sc-15383, RRID:AB_2071995 |
| Donkey anti-Mouse IgG (H+L) Highly Cross-Adsorbed Secondary Antibody, Alexa Fluor 488 | ThermoFisher | Cat# A-21202 |
| Donkey anti-Rabbit IgG (H+L) Highly Cross-Adsorbed Secondary Antibody, Alexa Fluor 488 | ThermoFisher | Cat# A-21206 |
| Donkey anti-Rat IgG (H+L) Highly Cross-Adsorbed Secondary Antibody, Alexa Fluor 594 | ThermoFisher | Cat# A-21209 |
| FLAG M2 | Sigma-Aldrich | Cat# F1804; RRID: AB_262044 |
| HA | BioLegend | Cat# 901502, RRID:AB_2565007 |
| HIS | Millipore-Sigma | Cat# H1029, RRID:AB_260015 |
| IRDye® 680RD Donkey anti-Mouse IgG Secondary Antibody | LI-COR Biosciences | Cat# 926-68072 |
| IRDye® 680RD Donkey anti-Rabbit IgG Secondary Antibody | LI-COR Biosciences | Cat# 926-68073 |
| IRDye® 680RD Goat anti-Rat IgG Secondary Antibody | LI-COR Biosciences | Cat# 926-68076 |
| IRDye® 800CW Donkey anti-Mouse IgG Secondary Antibody | LI-COR Biosciences | Cat# 926-32212 |
| IRDye® 800CW Donkey anti-Rabbit IgG Secondary Antibody | LI-COR Biosciences | Cat# 926-32213 |
| IRDye® 800CW Goat anti-Rat IgG Secondary Antibody | LI-COR Biosciences | Cat# 926-32219 |
| Myc | Santa Cruz | Cat# sc-40;H RRID: AB_627268 |

| | | |
|---|---|---|
| MYC | ThermoFisher | Cat# 13-2500, RRID:AB_2533008 |
| MYC | CCF Hybridoma Core | 9E10 |
| Phospho-Histone H3 S10 | CST | Cat# 3377; RRID: AB_1549592 |
| USP37 | Huang et al | N/A |
| B-ACTIN | Millipore-Sigma | Cat# A3854, RRID:AB_262011 |
| γH2AX | Millipore-Sigma | Cat# 16-193, RRID:AB_310795 |

**Bacterial and Virus Strains**

| | | |
|---|---|---|
| *Escherichia coli* BL21(DE3) Chemically Competent Cells | Thermo Fisher | Cat# C600003 |
| *Escherichia coli* NEB 5-alpha Chemically Competent Cells | NEB | Cat# C2987I |
| *Escherichia coli* NEB Stable Chemically Competent Cells | NEB | Cat# C3040I |

**Chemicals, Peptides, and Recombinant Proteins**

| | | |
|---|---|---|
| 5-Chloro-2′-deoxyuridine | MilliporeSigma | Cat# C6891 |
| 5-Iodo-2′-deoxyuridine | MilliporeSigma | Cat# I7125 |
| Acridine Orange | MilliporeSigma | Cat# A6014 |
| Aphidicolin | MilliporeSigma | Cat# 178273 |
| Cycloheximide | ThermoFisher | Cat# AC357420010 |
| Dynabeads | ThermoFisher | Cat# 10004D |
| Dynabeads Protein G | ThermoFisher | Cat# 10003D |
| Fluoromoun-G | ThermoFisher | Cat# 00-4958-02 |
| Glutathione Agarose Resin | MilliporeSigma | Cat# GE17-0756 |
| GST-CDH1 | This Paper | N/A |
| GST-USp37 | This Paper | N/A |
| HIS-Select Nickel Affinity Gel | MilliporeSigma | Cat# P6611 |
| Hydroxyurea | MilliporeSigma | Cat# H8627M |
| Lipofectamine RNAiMAX | Thermo Fisher | Cat# 13778150 |
| MG132 | R&D Systems | Cat# I-130 |
| Nocodazole | MilliporeSigma | Cat# M1404 |
| Odyssey® One-Color Protein Molecular Weight Marker | LI-COR Biosciences | Cat# 928-40000 |
| Paclitaxel | MilliporeSigma | Cat# 580556 |
| Pierce Anti-c-Myc Magnetic Beads | ThermoFisher | Cat# 88842 |
| Propidium Iodide | MilliporeSigma | Cat# P4864 |
| RNase A | MilliporeSigma | Cat# R6513 |
| Thymidine | MilliporeSigma | Cat# T1895 |
| TransIT-LT1 Reagent | Mirus Bio | Cat# MIR 2304 |

| | | |
|---|---|---|
| **Critical Commercial Assays** | | |
| Click-IT EdU Imaging Kit | ThermoFisher | Cat# C10340, C10639 |
| Pierce BCA Protein Assay Kit | ThermoFisher | Cat# 23225 |
| TnT® Quick Coupled Transcription/ Translation System | Promega | Cat# L2080 |
| **Experimental Models: Cell Lines** | | |
| 293T | ATCC | Cat# CRL-3216; RRID: CVCL_0063 |
| H1299 | ATCC (gift of A. Strohecker) | CRL-5803, RRID:CVCL_0060 |
| HCT116 | | |
| HeLa | ATCC | Cat# CCL-2; RRID: CVCL_0030 |
| MCF7 | ATCC | Cat# HTB-22; RRID: CVCL_0031 |
| MDA-MB-468 | ATCC | Cat# HTB-132; RRID: CVCL_0419 |
| T98G | ATCC | CRL-1690 , RRID:CVCL_0556 |
| U2OS | ATCC | HTB-96, RRID:CVCL_0042 |
| **Experimental Models: Organisms/Strains** | | |
| Zebrafish | | |
| **Oligonucleotides** | | |
| SMARTpool: ON-TARGETplus USP37 siRNA | Dharmacon | L-006085-00-0005 |
| ON-TARGETplus USP37 siRNA | Dharmacon | J-006085-06-0010 |
| ON-TARGETplus USP37 siRNA | Dharmacon | J-006085-07-0010 |
| ON-TARGETplus USP37 siRNA | Dharmacon | J-006085-09-0010 |
| Morpholino Oligo | GENETOOS, LLC | N/A |
| **Recombinant DNA** | | |
| pCDNA5-6xHIS-Ubiquitin | Burrows, et al. 2012 | |
| pCS2-FLAG | | |
| pCS2-MYC-AURORA B | This study | |
| pCS2-MYC-CHK1 | Pal, et al. 2019 | |
| pCS2-MYC-CHK1 L449R | Pal, et al. 2019 | |
| pCS2-MYC-USP37 | Burrows, et al. 2012 | |
| pCS2-TAP-USP37 | Huang, et al. 2011 | |
| pCS2-TAP-USP37 C350A | Huang, et al. 2011 | |
| pGEX6P1 | GE Lifesciences | Cat# 28954648 |
| pGEX6P1-CDH1 | Pal, et al. | |

| | | |
|---|---|---|
| pGEX6P1-USP37 | Burrows, et al. 2012 | |
| pLKO.1 shNonT | MilliporeSigma | Cat# SHC016 |

**Software and Algorithms**

| | | |
|---|---|---|
| FIJI | | https://imagej.net/Fiji |
| FlowJo X | FlowJo LLC | https://www.flowjo.com |
| Prism 7 | GraphPad | https://www.graphpad.com |
